# Unveiling drug tolerant and persister-like cells in *Leishmania braziliensis* lines derived from patients with cutaneous leishmaniasis

**DOI:** 10.1101/2023.07.04.547666

**Authors:** Marlene Jara, Jorge Arevalo, Alejandro Llanos, Frederik Van den Broeck, Malgorzata Anna Domagalska, Jean Claude Dujardin

## Abstract

**Introduction:** Resistance against anti-*Leishmania* drugs (DR) has been studied for years, giving important insights into long-term adaptations of these parasites to drugs, through genetic modifications. However, microorganisms can also survive lethal drug exposure by entering into temporary quiescence, a phenomenon called drug tolerance (DT), which is rather unexplored in *Leishmania*.

**Methods:** We studied a panel of 9 *Leishmania braziliensis* strains highly susceptible to potassium antimonyl tartrate (PAT), exposed promastigotes to lethal PAT pressure and compared several cellular and molecular parameters distinguishing DT from DR.

**Results and discussion:** We demonstrated *in vitro* that a variable proportion of cells remained viable, showing all the criteria of DT and not of DR: (i) signatures of quiescence, under drug pressure: reduced proliferation and significant decrease of rDNA transcription, (ii) reversibility of the phenotype: return to low IC_50_ after removal of drug pressure, (iii) absence of significant genetic differences between exposed and unexposed lineages of each strain and absence of reported markers of DR. We found different levels of quiescence and DT among the different *L. braziliensis* strains. We provide here a new *in vitro* model of drug-induced quiescence and DT in *Leishmania*. Research should be extended *in vivo*, but current model could be further exploited to support R&D, for instance, to guide screening of compounds able to overcome quiescence resilience of the parasite, hereby improving therapy of leishmaniasis.

## 1 Introduction

Chemotherapy is essential for the clinical management of infectious diseases; but also for their control and elimination, especially if humans constitute the reservoir of respective pathogens (anthroponoses) and drugs can counter human to human transmission (WHO, n.d.). Accordingly, treatment failure (TF) may have a major public health impact. TF is a clinical phenotype that can be expressed in different forms, like non-response, relapse or recrudescence. It has a complex and multifactorial origin, essentially involving the drug (quality, compliance, dosage), the host (immune status, co-infections and concomitant morbidities, existence of sanctuary organs/cells), and the pathogen’s biology (Barrett et al., 2019). From the microbial point of view, drug resistance (DR) is the usual suspect and culprit. Hence, DR is often and wrongly confounded with TF, while -in contrast with TF-drug resistance is a long-term parasite adaptive phenotype, more specifically a decreased drug susceptibility which is acquired through genetic modification of the microbe. As such, DR is heritable and the phenotypic adaptation can persist even in the absence of drug exposure (Balaban et al., 2019), until a mutation reverting or compensating the phenotype emerges and is selected.

However, other mechanisms can cause a reduction of drug-susceptibility in pathogens. Drug tolerance (DT) is one of them and it is intrinsically totally different from DR. Indeed, it refers to a short-term adaptive phenotype, consisting in decreased susceptibility of the pathogen to the drug, which is not due to an acquired genetic modification of the parasite. It is often associated with quiescence (syn. dormancy), a physiological state of the cell triggered by environmental insults and involving a reversible cell division arrest driven by a dynamic and regulated cell and metabolic remodeling program (Balaban et al., 2019). In microbiology, there are several examples of pathogens that enter in a transient quiescent state, in which they are refractory to one of more drugs, allowing them to persist in the host for long periods: among many others, *Mycobacterium tuberculosis*/Streptomycin, Isoniazid, Ciprofloxacin and Rifampin (Keren et al., 2011), *Staphylococcus aureus*/Aminoglycoside (Michiels et al., 2016) and *Plasmodium falciparum*/artemisinin (Teuscher et al., 2012).

*Leishmania* are parasitic protozoa causing a spectrum of clinical forms in (sub-)tropical regions but also in Southern Europe. Leishmaniasis belongs to the category of most neglected diseases, which is reflected among others by the small chemotherapeutic arsenal to combat them. Failure of the few available drugs jeopardizes elimination programs and some studies associated TF of leishmaniasis with DR (Lira et al., 1999). However, in two epidemiological settings (Peru/*L. braziliensis* and Nepal/*L. donovani*), we documented a poor correlation between resistance to antimonials (as measured by *in vitro* susceptibility assays) and TF (Yardley et al., 2006)(Rijal et al., 2007). This motivated us to look for alternative mechanisms leading to reduced drug susceptibility in the parasite, more specifically, quiescence. Research on quiescence in *Leishmania* is still in its infancy. Quiescence was reported to occur *in vitro* and *in vivo*, it is characterized by a reduced level of translational machinery (ribosomal RNA and proteins), low rates of RNA synthesis, protein turnover and membrane lipid synthesis (Kloehn et al., 2015). Last but not least, parasite populations are found to be heterogeneous in terms of activity, with deeply and semi-quiescent subpopulations (Mandell and Beverley, 2017).

There are only two studies on *Leishmania* quiescence in the context of drug exposure. First, *in vivo*, following treatment with Miltefosine, quiescent cells of *L. mexicana* were encountered in mesothelium-like tissues surrounding granulomas, in which the drug did not accumulate (Kloehn et al., 2021). Secondly, *in vitro*, we showed that a strain of *L. lainsoni* entered in quiescence under stationary phase or antimony pressure. Several transcriptional signatures were shared by the parasites surviving to both environmental insults: among these, a series of transcripts were present in higher abundance, in a general background of transcriptional shift (Jara et al., 2022). These molecular similarities validated the quiescent character of *Leishmania* under antimony pressure. The objective of the present study was to further explore -*in vitro*-the link between quiescence and DT in *Leishmania* and to address the diversity of the DT phenotype in genetically different strains of the same species. Therefore, we exposed 9 strains of *L. braziliensis* to high doses of trivalent antimonials and compared several parameters distinguishing DT from DR in pre- and post-exposure lineages (Inhibitory concentration, proliferation, cell viability, metabolic activity, lethal dose, and genome).

## 2 MATERIALS AND METHODS

### 2.1 Parasites

Clinical isolates were collected in Peru at the Institute of Tropical Medicine Alexander von Humboldt between 2001-2004 within the framework of the multi-regional LeishNatDrug project (Yardley et al., 2006). Nine of them were selected for present study, according to the following inclusion criteria: (i) first episode of leishmaniasis, (ii) cutaneous form of the disease, (iii) no previous treatment before attending to the clinic, (iv) complete therapy with antimonial post-diagnosis, (v) no concomitant disease and (vi) infecting species typed as *Leishmania braziliensis*. One strain of *Leishmania lainsoni* was included in the study as control: this is a line highly susceptible to antimonials, for which the existence of quiescence after PAT exposure was demonstrated (Jara et al., 2022). For each isolate, enhanced green fluorescent protein (EGFP) was integrated within the 18S ribosomal DNA locus, further called rEGFP with the use of the pLEXSY-hyg system (Jena Bioscience) as previously reported elsewhere (Bolhassani et al., 2011; Jara et al., 2022). After the generation of the transgenic rEGFP parasites, clones were obtained with the ‘micro-drop’ method, as described elsewhere (Van Meirvenne et al., 1975; Jara et al., 2022). All the work presented here was made with these clones, further called rEGFP strains.

### 2.2 Cell culture of promastigotes and generation of axenic amastigotes

Promastigotes were maintained in complete M199 (M199 medium at pH 7.2 supplemented with 20% fetal bovine serum, hemin 5 mg/L, 50 μg/mL of hygromycin Gold, 100 units/mL of penicillin and 100 μg/mL of streptomycin) at 26 °C with passages done twice per week. To obtain axenic amastigotes, 1 mL of stationary promastigotes was centrifuged (1500 g, 5 min), the pellet was re-suspended in 5 mL of complete MAA (M199 at pH 5.5, supplemented with 20% fetal bovine serum, glucose 2.5 g/L, 5 g of tryptic soy broth, hemin 5 mg/L, 25 μg/mL of hygromycin Gold, 100 units/mL of penicillin and 100 μg/mL of streptomycin) and parasites were incubated at 34 °C. After observing morphological evidence of amastigogenesis at day 3, the lines were subcultured each 4 days and growth was monitored microscopically over three weeks. If axenic amastigotes were not able to proliferate, three subsequent attempts for axenization were performed: the generation of axenic amastigotes was successful for 3 rEGFP strains (PER094, 122 and 362). Samples for further experimental procedures were prepared in complete M199 or MAA in absence of hygromycin. The growth curves were monitored by daily counting of parasites and the generation time was calculated with the formula G=t/ (logb-logB)/log2). Where t = time interval in hours, B = number of parasites at the beginning of a time interval, b = number of parasites at the end of the time interval.

### 2.3 Cell recovering and monitoring of survival after drug pressure

Exponentially growing promastigotes were exposed to 9 µg/mL of potassium antimonyl tartrate (PAT) for 48 hrs after which an aliquot of 50 µL was sub cultured in a medium without drug pressure (Post-PAT). For each strain, a control without exposure to drug pressure and instead treated with PBS was maintained in parallel (Post-PBS). The cell cultures were monitored microscopically, and if positive cellular growth was observed, additional subcultures for the evaluation of the drug susceptibility and pellets for sequencing were prepared. Throughout the paper, lineages that were exposed to PAT were referred to as ‘Post-PAT’, while those that were not previously exposed to PAT were referred to as ‘Post-PBS’.

### 2.4 Drug susceptibility and estimation of the IC_50_

The drug susceptibility was measured with the resazurin test. Briefly, exponentially growing parasites were plated into 96 wells plates containing a serial dilution of PAT to reach final concentrations ranging from 83 to 0.1 µg/ mL. Each plate included controls without PAT and controls for monitoring proliferation and autofluorescence of the medium. Resazurin sodium salt (200 µg/mL, Sigma) was added at two different time points: at the time of plating for the proliferation controls (T1) and 20 hrs post drug pressure for the other wells (T2). After 4 hrs of incubation with resazurin, the fluorescence of its reduced form resorufin was recorded using the Victor X3 Multilabel Reader (PerkinElmer) exciting at 560 nm and measuring emission at 590 nm. The test was considered valid for further analysis only if the fluorescence for the controls without drug pressure at T2 was at least 2 fold the fluorescence for the proliferation controls at T1. All experiments with the different lines were repeated 3 times with three technical replicates each. The blank-subtracted data expressed in relative fluorescence units (RFU) were exported to GraphPad Prism 8 to calculate the 50% inhibitory concentration (IC_50_), using a sigmoidal dose-response model with variable slope. Statistical analysis and data visualization were performed in R studio with built-in functions and ggplot2, respectively.

### 2.5 Flow cytometry assay for monitoring single cell viability and quiescence

Exponentially proliferating cells (promastigotes and amastigotes) were expose to PAT at concentration ranging from 1 to 83 µg/ mL and both the cell viability and the rEGFP expression were measured by flow cytometry at three time points: before drug exposure and after drug pressure at 24 and 48 hrs. The rEGFP expression is a negative marker of quiescence: highly expressed in proliferative cells and downregulated during quiescence (Jara et al., 2019, 2022). The cell viability was evaluated by preincubating cells with the NucRed dead 647 (Thermo fisher scientific) for the staining of dead cells and Vybrant Dye Cycle violet (Thermo fisher scientific) for the staining of DNA in all cells. The Vybrant Dye Cycle violet was used to select the subpopulation of cells having a healthy pattern for their DNA. A wild type (non rEGFP) strain and a non-stained sample were included as negative controls, together with a sample exposed to thermal shock as a positive control for cellular death. The samples were analyzed with a calibrated flow cytometer BD FACS Verse ™ in the medium flow rate mode. In order to compare the rEGFP relative fluorescence units (RFU) among samples, the acquisitions were made with the same settings during all the experiments. The FCS files were analyzed with the *FCS 5 express Plus research* edition. The subpopulation of cells with good cell viability was selected by sequential gating. Briefly, single cells were selected by pulse geometry gate; a first gate was selected by plotting the SSC-W vs SSC-H, and a second gate was created by plotting the FCS-H and FCS-A. Among single cells, a third gate was created by plotting the rEGFP vs the NucRed and selecting the subpopulation NucRed-negative. Finally a fourth gate was created by plotting the rEGFP vs the Vybrant Dye Cycle violet and selecting for the subpopulation having a healthy DNA pattern. The cell viability for each sample was estimated by multiplying the percentage of cells NucRed-negative and Vybrant DyeCycle-positive. For each developmental stage of *Leishmania* and for each time point post exposure to PAT, the effect of the drug pressure over the rEGFP expression in the subpopulation of viable cells was evaluated with a two-way ANOVA considering the PAT drug pressure and strains as explanatory variables. Subsequently a Tukeys HSD test was used to evaluate the statistical significance of differences between pairwise comparisons.

### 2.6 Estimating the lethal dose 10, 20 and 50

The measures of cell viability obtained after the flow cytometry assay over increasing concentration of PAT were used to estimate the lethal doses of PAT that kill 10, 25 and 50 % of the population (LD_10_-LD_50_). The LDs were estimated using the *drc* R package(Ritz et al., 2015), briefly: the data were fitted using a log-logistic model with a lower limit at 0 and having as formula the % of cell viability as explained by the concentration of PAT and the line of the parasite as grouping factor. Finally, the desired LDs were estimated having as reference the upper limit and setting the confidence interval to 0.95.

### 2.7 Whole genome sequencing

For each lineage (post-PBS and post-PAT), DNA was isolated from a sample maintained without drug pressure and a sample recovered after 48 hrs of PAT drug pressure, using the QIAamp DNA Micro Kit (Qiagen). The DNA concentration was assessed with the Qubit DNA broad-range DNA quantification kit (Thermo Fisher). Library preparation and sequencing were performed at BGI with a DNBSEQ-WGS-PCR free library index and the DNBSEQ PE150 platform, respectively. The FASTQ files containing paired reads of 150 bp were subsequently analysed for the assessment of sequence variants and ploidy. Briefly, quality of the reads was evaluated with samtools and the multiqc command. Samples were aligned against the reference genome of *L. braziliensis* MHOM/BR/75/M2904 (Van den Broeck et al., 2020). The per position mean depth and the breath of coverage were estimated with SAMtools. Variants (SNPs and INDELs) were called jointly for all 18 samples with BCFtools and the mpileup command (Danecek et al., 2021). Subsequently, the multisamples vcf file was filtered according to following criteria: overall variant quality (QUAL) higher than 200, the individual read depth per locus (FMT/DP) higher than 5, and the average mapping quality (MQ) and genotype quality (GQ) higher than 40. The annotation of the genomic location for the variants was done with SnpEff (Cingolani et al., 2012). Somy values were estimated by using the median read depth across each chromosome as reported elsewhere (Dumetz et al., 2017). A maximum likelihood phylogenetic tree was reconstructed with IQTREE v1.6.12 (Nguyen et al., 2015) reusing HKY+G substitution model and 1000 ultrafast bootstraps approximation (Hoang et al., 2018).

## 3 Results

### 3.1 *Leishmania* exposure to transient PAT drug pressure causes a reversible decrease of proliferation and does not select for parasites with lower drug susceptibility

For each of the 9 rEGFP strains of *L. braziliensis* used, we measured the drug susceptibility of promastigotes of the corresponding post-PBS lineages), with a resazurin test. Values ranged from 0.2 to 7.48 µg/mL, with a median of 1.98 µg/mL (**Figure 1A** and **Supplementary Figure 1**). We then exposed all lines to the same concentration of PAT (9 µg/mL, representing ∼4 fold the median IC_50_ of all 9 strains) in order to evaluate potential differences in their adaptation to survive. (i) The potential cytostatic effect of PAT on *Leishmania* cells, was monitored with the resazurin test by measuring the signal of its reduced form resorufin before the drug treatment and after 24 hrs of exposure to PAT or PBS. In the absence of the drug, the resorufin log2 FC signal ranged from 0.8 (PER206 EGFP Cl2) to ∼ 3.2 relative fluorescence units (RFU, equivalent to 9 fold the initial signal, in PER122 EGFP Cl1), indicating that all strains were proliferating albeit at different rates. Under drug pressure, the resorufin log2 FC signal was significantly decreased compared to the one measured in the corresponding control without the drug in all strains suggesting that under our conditions of drug pressure, parasites have very diminished growth (**Figure 1B**). (ii) After 48 hrs of exposure to 9 µg/mL of PAT, parasite’s death was observed in each lineage, but parasites were all able to resume their proliferation after subculturing in a fresh medium without drug. This indicated that at least a proportion of the cells survived and maintained their capability to switch to a proliferative state once they are in optimal conditions. (iii) We found that the susceptibility to the drug did not change in the post-PAT lineages (median IC_50_ = 2.35 µg/mL; interquartile range, IQR= 2.54) when compared to the corresponding post-PBS lineages (median IC_50_ = 2.00 µg/mL, IQR= 2.70, Mann–Whitney U test, P= 0.9, **Figure 1A**). Noteworthy, the capacity of *Leishmania* to survive concentrations of drug beyond the IC_50_ of 9 µg/mL represents ∼ 45 times the IC_50_ for the most susceptible line PER348 EGFP Cl1 (IC50= 0.2 µg/mL).

**Figure 1.**
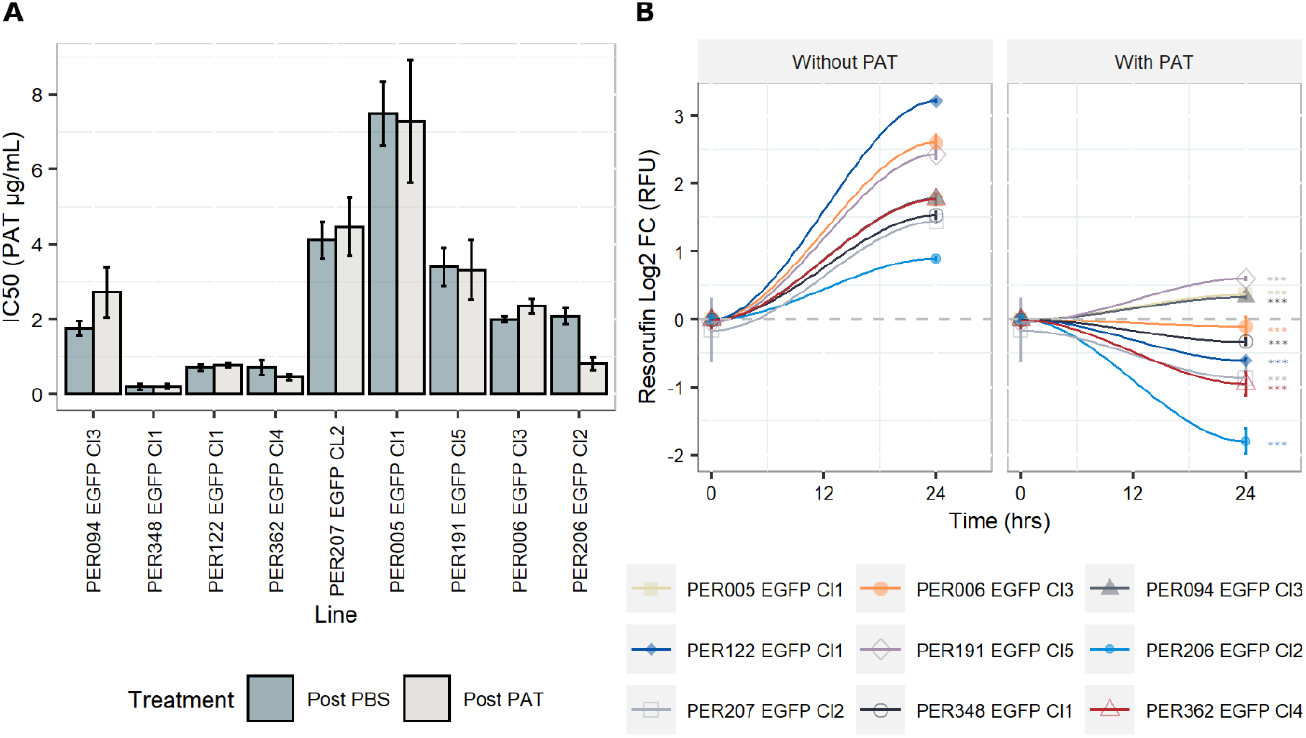
Drug susceptibility of *L. braziliensis* promastigotes and their growth features under drug pressure. **A**. Drug susceptibility to PAT in 9 strains as estimated by the resazurin test after 24 hrs of drug pressure. For each strain, the IC_50_ was calculated in a lineage without prior exposure to the drug (Post-PBS) and after the exposure to 9 µg/ml of PAT (∼ 4 fold the median IC50 considering all lineages, Post-PAT). The results represent the mean ± SEM of three biological replicates. **B**. Evaluation of the cytostatic effect of PAT at 9 µg/ml. The gray dashed line represents no change in the original resorufin signal compared to the day the cells were initially plated. A Log2 FC above 1 indicates parasite duplication and a Log2 FC below 0 indicates parasites arrest and/or cellular death. The asterisks represent statistically significant differences after Tukey’s post-hoc test (p < 0.001) between cells after 24 hrs of plating without drug compared to cells under drug pressure.

### 3.2 *Leishmania* exposure to transient PAT drug pressure is accompanied by a decrease in the expression of rEGFP among viable cells

The expression of rEGFP is a negative biomarker of quiescence: it is high in proliferative cells while it decreases or disappears in quiescent cells (Jara et al., 2022). (i) In promastigotes, the exposure of proliferating cells to PAT (9.0 µg/mL) led to a significant reduction in the levels of rEGFP expression (RFU) among viable cells of each line when compared to the control treated with PBS at both 24 and 48 hrs (**Figure 2A**). Furthermore, rEGFP expression was also different among the evaluated strains (two-way ANOVA: P <0.001 for PAT treatment and P<0.001 for strain). The same significant effect of the drug was observed in amastigotes. (ii) We then explored if the reduction in rEGFP expression changes in response to increasing concentrations of PAT. The results showed that higher doses of PAT induced a more profound reduction of the rEGFP expression compared to the control without PAT treatment at both 24 hrs and 48 hrs post PAT treatment in both promastigotes and amastigotes (**Figure 2B**). The reduction in rEGFP expression follows a dose-response model where there is a linear response that reaches a plateau around 9.0 µg/mL of PAT with an average relative rEGFP reduction of 44 and 40 % after 48 hrs of PAT in promastigotes and amastigotes respectively. (iii) The analysis of relative rEGFP reduction also shows differences among *L. braziliensis* strains (ranging between 23.7 and 56.2 % in promastigotes and between 33.7 and 49.9 % in amastigotes).

**Figure 2.**
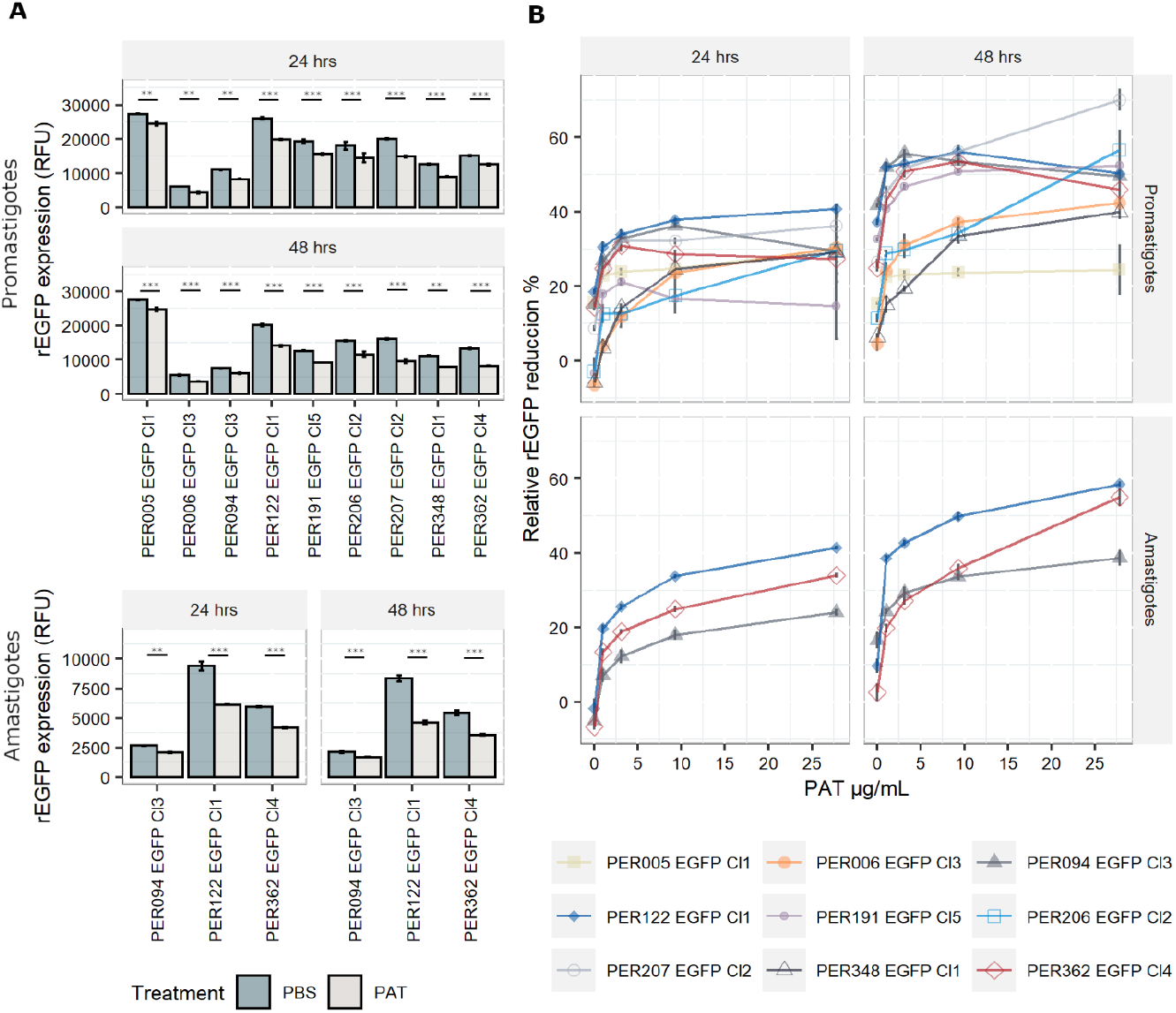
rEGFP expression in promastigotes (9 strains) and amastigotes (3 strains) of *L. braziliensis* under drug pressure; measures are made on viable cells. **A**. rEGFP expression of *L. braziliensis* lineages after 24 and 48 hrs of exposure to 9 µg/mL of PAT. The results represent the mean ± SEM of three biological replicates. The asterisks represent statistically significant differences after Tukey’s post-hoc test; ^**^ *p* < 0.01, ^***^ *p* < 0.001. **B**. Relative rEGFP reduction (in %) in relationship to increasing concentrations of PAT after 24 and 48 hrs of PAT exposure.

### 3.3 Survival of Leishmania to high lethal doses of PAT is accompanied by decrease in rEGFP expression

When measuring the drug susceptibility with standard fluorometric procedures such as the resazurin test at a single point of time, it is not possible to address with certainty if the drug has cytostatic or cytotoxic effects. The decreased signal of resorufin in samples with the drug pressure compared to the control could result from cellular death, arrested growth, or likely, a mixture of both events. Therefore, we evaluated the cell viability at the single-cell level by flow cytometry and calculated the lethal doses 10, 25, and 50 (LD_10_, LD_25_, and LD_50_), which are the concentrations of drug that kill 10, 25, and 50% of the population respectively. (i) In promastigotes, the kinetics of the cell viability over different concentrations of PAT pressure suggests variable cytotoxic effects among the evaluated lines. The LD_50_ among all strains ranged from 5.98 µg/mL in PER206 EGFP Cl2 to 83.3 µg/mL in PER006 EGFP Cl3 (**Figures 3A-B**). Median LD_50_ among the 9 strains was 28.2 µg/mL being about 10 fold the IC_50_ that was previously measured with the resazurin test. Noteworthy 4 out of the 9 lines had LD_50_ higher than 30 µg/mL of PAT, a non-physiological concentration. These strains may be considered as highly tolerant to PAT compared to the other strains at the same parasite stage. (ii) As promastigotes showed high variability in their rates of survival to increasing concentrations of PAT, we further evaluated the potential relationship between the capability to adopt a quiescent state as measured by the reduction in the rEGFP expression and the success to overcome the drug pressure as measured by their cell viability. After 24 hrs of PAT treatment there was a weak positive relationship at low concentrations of PAT (1 to 3 µg/mL of PAT) that became strong at 9 and 28 µg/mL of PAT (**Figure 3C**). (iii) Amastigotes showed remarkably higher survival rates to PAT compared to the promastigotes of the respective strains. After twice the time of exposure to PAT (48 hrs) the highest concentration of the drug could not kill 50% of the population, and the estimation of the median LD_25_ for the three strains was 46.7 µg/mL (**Figure 3D-E)**. Because these results with amastigotes of *L. braziliensis* were surprisingly high, we evaluated the LDs from a highly susceptible line of *L. lainsoni*. The same experimental setup estimated that in amastigotes the LD_25_ and LD_50_ were 0.49 and 0.97 µg/mL respectively, ruling out that the *L. braziliensis* results would be an experimental artifact (**Figure S2**). Accordingly, amastigotes of *L. braziliensis* are intrinsically more tolerant to PAT pressure compared to promastigotes.

**Figure 3.**
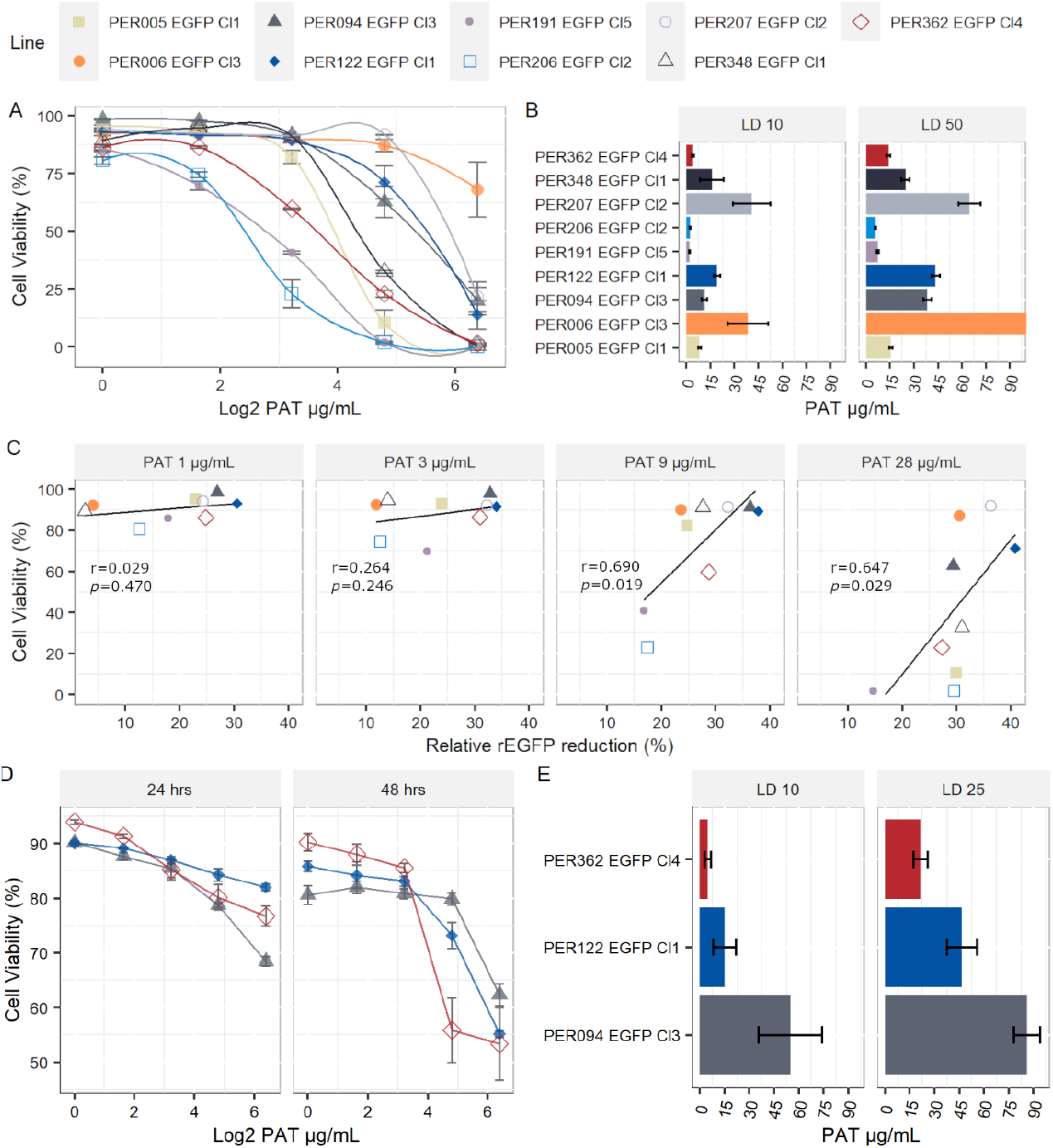
Lethal dosis (LD) on *L. braziliensis* promastigotes and amastigotes. **A**. Viability of individual promastigotes after 24 hrs of exposure to increasing PAT concentration. **B**. Lethal doses 10 and 50 on promastigotes. The results represent the mean ± SEM of three biological replicates. **C**. Relationship between viability of promastigotes and reduction of rEGFP expression at different PAT concentrations. The Pearson’s correlation coefficients and p values are shown. **D**. Viability of individual amastigotes after 24 and 48 hrs of exposure to increasing PAT concentration. **E**. Lethal doses 10 and 25 on amastigotes, calculated after 48 hrs of PAT exposure. The results represent the mean ± SEM of three biological replicates.

### 3.4 Leishmania lines exposed to PAT are genomically similar to their respective parental lines

We evaluated the genome stability between pre- and post-PAT lines by WGS, at the levels of somy, SNPs and indels. (i) At somy level, we observed variability of karyotypes in a strain-specific manner. Number of aneuploid chromosomes/strain ranged from 1 to 10. Trisomy of chromosomes 5, 11, 25, 29, and tetrasomy of chromosome 31 were among the most frequent occurrences being present in at least 3 out of the 9 strains. For each strain, the original overall ploidy of the post-PAT lineage was identical to the corresponding post-PBS lineage (**Figure. 4A**). (ii) Compared to the *L. braziliensis* reference genome (MHOM/BR/75/M2904), a total of 163,823 high-quality SNPs and 17,055 high-quality INDELs were called across our panel of 18 lineages. A maximum likelihood phylogenetic tree based on SNPs shows that each post-PAT lineage clusters with its corresponding control, post-PBS lineage (**Figure 4B**), reflecting the high genomic similarity between post-PAT and post-PBS lineages. Indeed, each post-PAT lineage showed differences to its corresponding control lineage at only 11 to 208 SNP loci and 76 to 119 INDEL loci, the majority of which (73.4% SNPs and 93.1% INDELs) occurred in the noncoding region of the genome (**Supp. data**). None of these variants occurred in a homozygous state; in other words, all variants were either absent in the post-PBS lineage and heterozygous in the post-PAT lineage, or vice versa. The genomic analysis indicates -for each strain-that PAT-exposed lines are genomically very similar to their parental line and that although a limited number of mutations occur they are not in genes known to be related with drug resistance in *Leishmania*.

**Figure 4.**
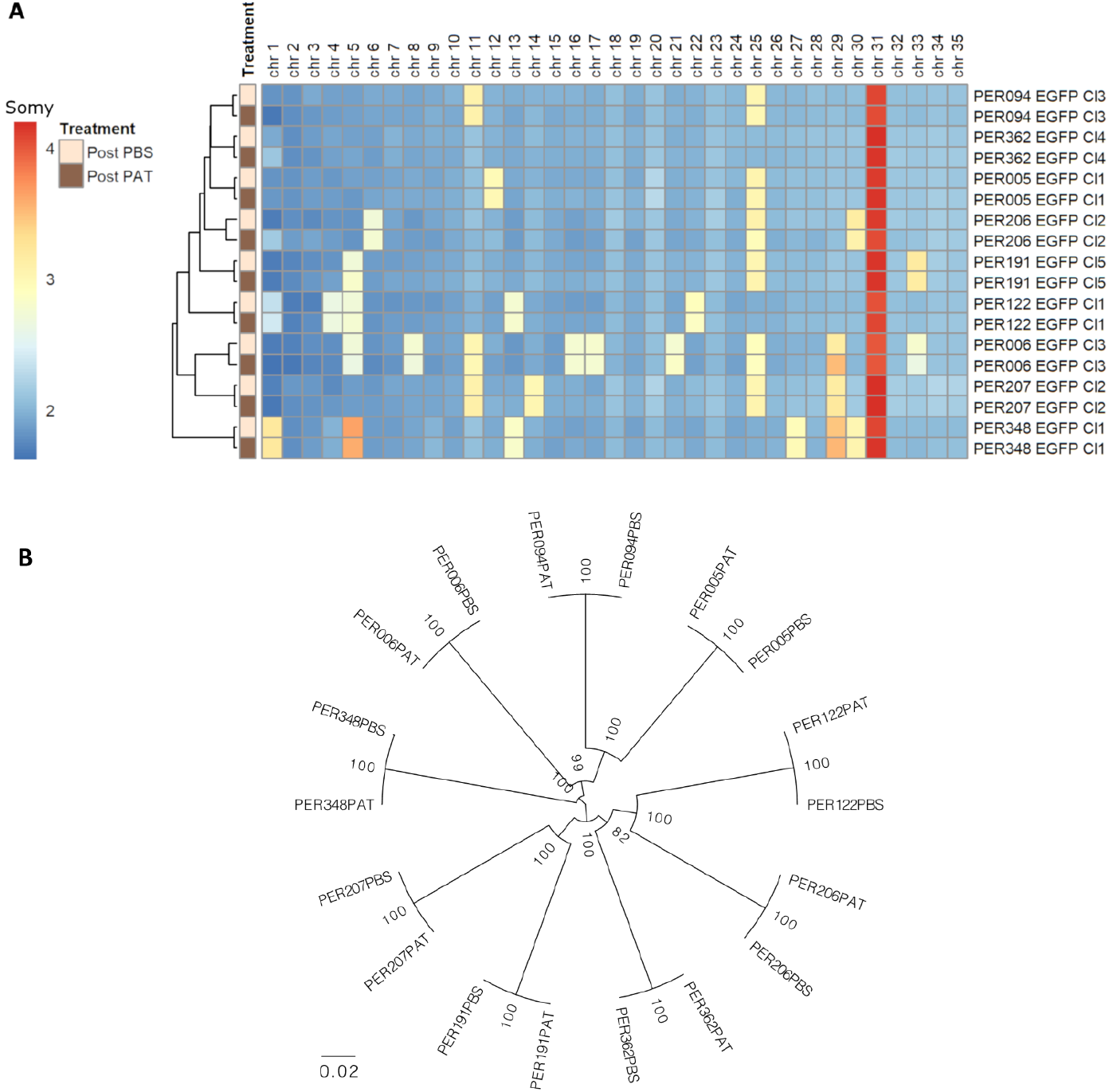
Genomic differences between the 9 strains of *L. braziliensis* and each lineage before (Post-PBS) and after (Post-PAT) drug pressure; bulk whole genome sequencing. **A**. Ploidy. **B**. maximum likelihood phylogenetic tree based on 163,823 high-quality SNPs.

### 3.5 After longer and stronger exposure to environmental stress, a small fraction of cells survive and show a reduced rEGFP expression

Persisters is another phenotype associated with quiescence (Barrett et al., 2019) and they usually represent a very small fraction of the population. We explored the occurrence of persister-like cells in promastigotes under two conditions: We explored the survival of cells under two conditions of stronger stress: by long-term stationary cuture of PER191 EGFP Cl5, where cells were maintained without subculturing over 14 days (Sta D14) and by exposing exponentially growing cells to PAT and maintaining the drug pressure over 14 days (Log PAT D14). We tested the resilience of the survivors by treating them with an exposure to PAT or PBS as untreated control over 24 hrs (Fig. 5 A). (i) On one hand, the Sta D14 population treated with PBS showed an average cell viability of ∼30% [3 replicates, only one showed in Fig.5 B] while cells treated with PAT showed a significant decrease of their average viability to ∼3.56% (T test, *p*<0.001). On the other hand, the Log PAT D14 population showed a very small fraction of survivors (average of ∼0.63%) and after a second exposure to PAT, there was no significant change in the proportion of survivors (average of ∼0.95%)). (ii) There was a significantly higher reduction of the levels of rEGFP expression compared to what was initially observed in cells exposed to 48 hours of PAT (Fig.2): 87.5 % vs a maximum of 56.2 %, respectively. There was no further rEGFP reduction when Sta D14 and Log PAT D14 samples were (further) exposed to PAT, suggesting that the minimum basal levels of expression required for survival were reached. (**Figure 5 A-B**). Besides the analysis by flow cytometry, we also looked for the presence of survivors under the confocal microscope. It was possible to observe a very scarce number of cells that had very low levels of rEGFP expression and that despite remaining at the same location had a very clear motility of their flagellum. Survivors from both conditions were able to proliferate after passing them in a fresh medium. The new population reached 95 and 85 % of cell viability by day 7 in Sta D14 and Log PAT D14 respectively.

**Figure 5.**
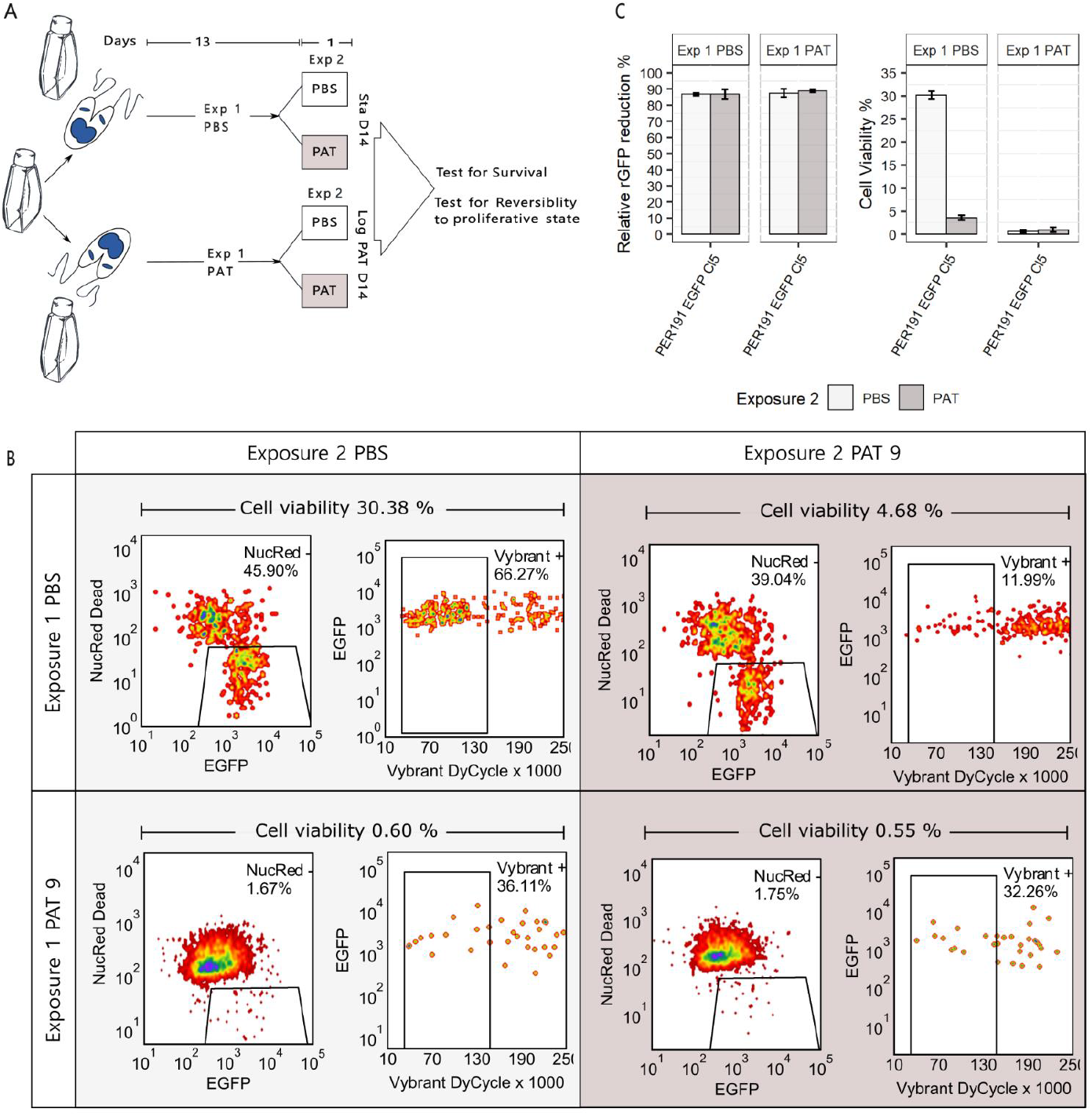
Promastigote (PER191 EGFP Cl5) viability under conditions of longer and stronger environmental stress. A. Experimental outline. Cell viability as measured by flow cytometry in (i) long term stationary phase, at day 14 (Sta D14) or (ii) after exposing proliferating cells to 14 days of PAT pressure (Log PAT D14); we tested the resilience of the survivors on day 14 after treating them with a second exposure to PAT or PBS as control over 24 hrs. The concentration of the first and second exposure to PAT was 9 µg/mL. See Materials and Methods for details on sequential gating. B. The density plots showing the percentage of survivors on day 14 in one representative biological replicate per experimental condition. C. Cell viability and their levels of relative rEGFP reduction in long-term cultures as described in panel A. The bars represent the mean ± SEM of three biological replicates. The asterisk represents statistically significant differences after a T test, ^***^ *p* < 0.001.

## 4 Discussion

We studied here the resilience of *L. braziliensis* cells surviving to high doses of PAT *in vitro*. Exposure to transient PAT pressure caused a reversible decrease of cell proliferation as well as a significant drop of rDNA transcription, measured here by a decrease of the rEGFP expression among viable cells. The coincidence of these 2 features is a hallmark of quiescence under stressful conditions (Jara et al., 2022), but was also observed in a *L. lainsoni* strain exposed to PAT (Jara et al., 2022). We also found in present study that PAT exposure did not select for parasites with lower drug susceptibility and that for each *L. braziliensis* strain, pre- and post-PAT lines were genomically similar and no markers of DR were encountered. Altogether, these 4 observations suggest that surviving PAT exposure in our experimental conditions was not due to DR, well to a form of quiescence-mediated DT.

Quiescence is a widespread adaptive strategy which allows cells to survive environmental insults. Pathogenic vector borne parasites like *Leishmania* are ‘naturally’ exposed to multiple stresses in the insect vector and the mammal host (Caljon et al., 2016) and quiescence could play a major adaptive role among other survival skills. Exposure to drugs is evolutionary recent an constitutes and ‘artificial’ source of stress to which parasites can adapt by genetic variation (DR) or by the exploitation of pre-existing mechanisms of quiescence, through metabolic modulation (DT) as demonstrated here, with PAT. Under high concentrations of PAT (median LD_50_ of 28 µg/ml), a large proportion of cells remained viable and this was significantly correlated with decrease in metabolic activity as measured by the relative rEGFP reduction. As such, this is close to the definition of DT as given in bacteriology: *i.e*. the general ability of a population of cells to survive longer treatments, a.o. by having a lower killing rate (Balaban et al., 2019), here measured by the LD_50_ which was about 10 times the IC_50_. Pushing our experimental conditions further -by longer exposure to environmental stress: long term stationary phase and/or PAT exposure-showed that there were still survivors, albeit a smaller proportion than under short exposure to drug. This was accompanied by a higher reduction of the rEGFP expression, indicating a deeper quiescent stage in those survivors. These highly resilient parasites are very similar to persisters, a subpopulation of tolerant cells in bacteria, which can survive antibiotic treatment much better than the majority of the population (Balaban et al., 2019).

The study was done with axenic promastigotes (model for the developmental life stage of the sand fly) and results were confirmed in axenized amastigotes (models for the developmental life stage living in the mammal). Further work is required to extend and confirm these results in an intracellular context and in vivo. The only report currently available on in vivo quiescence in the context of chemotherapy revealed, in Miltefosine-treated *L. mexicana*-infected BALB/c mice, the presence of quiescent amastigotes in collagen-rich, dermal mesothelium surrounding granulomas (Kloehn et al., 2021): authors concluded that quiescence, together with the lack of miltefosine accumulation in the mesothelium may contribute to drug failure and nonsterile cure. The potential for DT and persistence shown here by *L. braziliensis* strains and by *L. lainsoni* (Jara et al., 2022) might explain several (sub-)clinical features associated with species of subgenus *Viannia*, like (i) the presence of parasites in 80% of scars, years after treatment with pentavalent antimony therapy (Schubach et al., 1998); (ii) the mucosal metastases from healed primary cutaneous lesions (Jha et al., 2022) and (iii) the treatment failure in absence of DR (Yardley et al., 2006). Longitudinal studies of infected individuals and monitoring of quiescence is needed to test these hypotheses. Such studies are not likely to be easy given the paucity -by definition-of persister cells and they require positive markers (and not negative markers, like the rEGFP that can be used in experimental studies like here). A first step in that direction was made by a transcriptomic study which indicated that in an overall context of transcriptional shift, some transcripts were relatively more abundant in quiescent cells, like leishmanolysin (GP63), amastin and amastin-like proteins, and autophagy-related genes (Jara et al., 2022).

Noteworthy, the 9 *L. braziliensis* strains here studied showed different levels of quiescence and DT, which was not associated with the large genetic distances between each of the 9 strains: for instance, the 4 strains with the highest LD_50_ (PER006, 094, 122 and 207) were spread over the phylogenetic tree shown in Fig.4B. There was also no apparent link with the treatment outcome of the patients from which the parasites were originating: for instance, PER006 and PER206, which showed respectively the highest and lowest values of LD_50_ both came from unresponsive lesions. However, it is too premature to infer any impact of DT level measured in vitro on the treatment outcome in patients. Indeed, a huge selection bias is introduced when isolating and cultivating parasites. There is a strong bottleneck at isolation time and during in vitro maintenance, the parasites which are fittest in the culture medium will be selected and will form a dominant population; this is not necessarily the same as the dominant population present in the patient, a phenomenon that we observed in *L. donovani* (Domagalska et al., 2019). Accordingly, in a next step, quiescence and DT should -as previously recommended for DR (Domagalska and Dujardin, 2020)-also be studied directly in the patient to know its clinical impact and discriminatory molecular markers as well as sensitive detection methods should be developed for this.

It is very likely that the results here observed are drug-specific and cannot be generalized to all antileishmanial drugs. Indeed, quiescence is expected to protect against drugs (i) interfering with the parasite metabolism and (ii) which can be countered by decreasing this metabolism. This is the case for three of the anti-leishmania drugs: Pentavalent antimonials (Sb^V^), Miltefosine (MIL) and Paromomycin (PMM). The reduced form of Sb^V^ (PAT) has a direct effect on the parasite by disturbing its redox-potential (Wyllie et al., 2004) and a similar effect is expected with Sb^V^ itself which has an indirect effect through interfering with signaling of the macrophage and triggering ROI/RNI production in the host cell (Basu et al., 2006). MIL interferes with the biosynthesis of lipids including sterols, sphingolipids (Armitage et al., 2018) and the metabolism of alkyl-lipids, also inducing mitochondrial depolarization and a decrease of intracellular levels of ATP (Ponte-Sucre et al., 2017). PMM targets the decoding A-site of the ribosomes’ small subunit increasing misreading and translation inhibition (Chawla et al., 2011). In contrast, Amphotericin B which binds to ergosterol-related sterols in the cell membrane, inducing the production of a pore and fatal exchange of ions (Ramos et al., 1996) is expected to induce less (if any) quiescence and DT.

Research on quiescence, drug tolerance and persistence in *Leishmania* is still in its infancy. Besides providing a knowledge that might help better understanding the pathophysiology of *Leishmania* infections, it might also support and guide further investigations on new chemotherapeutic interventions to counter the disease as well as the infection. Historically, research into new drugs screened replicative forms, creating a leishmaniasis intervention tool kit that neglects the impact of quiescent forms. We provide here a new model of quiescence and DT that could be further exploited for in vitro screening of compounds able to overcome quiescence resilience of the parasite, hereby improving therapy of leishmaniasis.

## 5 Conflict of Interest

The authors declare that the research was conducted in the absence of any commercial or financial relationships that could be construed as a potential conflict of interest.

## 6 Author contributions

Substantial contributions to the conception or design of the work; or the acquisition, analysis, or interpretation of data for the work: all.

Drafting the work or revising it critically for important intellectual content: all

Provide approval for publication of the content: all

Agree to be accountable for all aspects of the work in ensuring that questions related to the accuracy or integrity of any part of the work are appropriately investigated and resolved: all

## 7 Funding

This study was financially supported by the Flemish Fund for Scientific Research (postdoctoral grant to MJ, FWO-1223420N and postdoctoral grant to FVdB, FWO-1226120N), the European Commission (contracts ICA4-CT-2001-10076 and INCO-CT2005-015407) and the Flemish Ministry of Science and Innovation [SecondaryResearch Funding ITM – SOFI, Grant MADLEI].

## 8 Acknowledgments

We acknowledge Ilse Maes for her logistic support in the lab.

## 10 Supplementary Material

See attached

## 11 Data Availability Statement

The raw data from the WGS for this study can be found in the Sequence Read Archive (SRA) submission: SUB12410472 (https://submit.ncbi.nlm.nih.gov/subs/sra/SUB12410472/overview).

## Notes

https://submit.ncbi.nlm.nih.gov/subs/sra/SUB12410472/overview

### Competing Interest Statement

The authors have declared no competing interest.

