## Supplemental Figures 1 and 2 for "Unveiling drug tolerant and persister-like cells in *Leishmania braziliensis* lines derived from patients with cutaneous leishmaniasis"

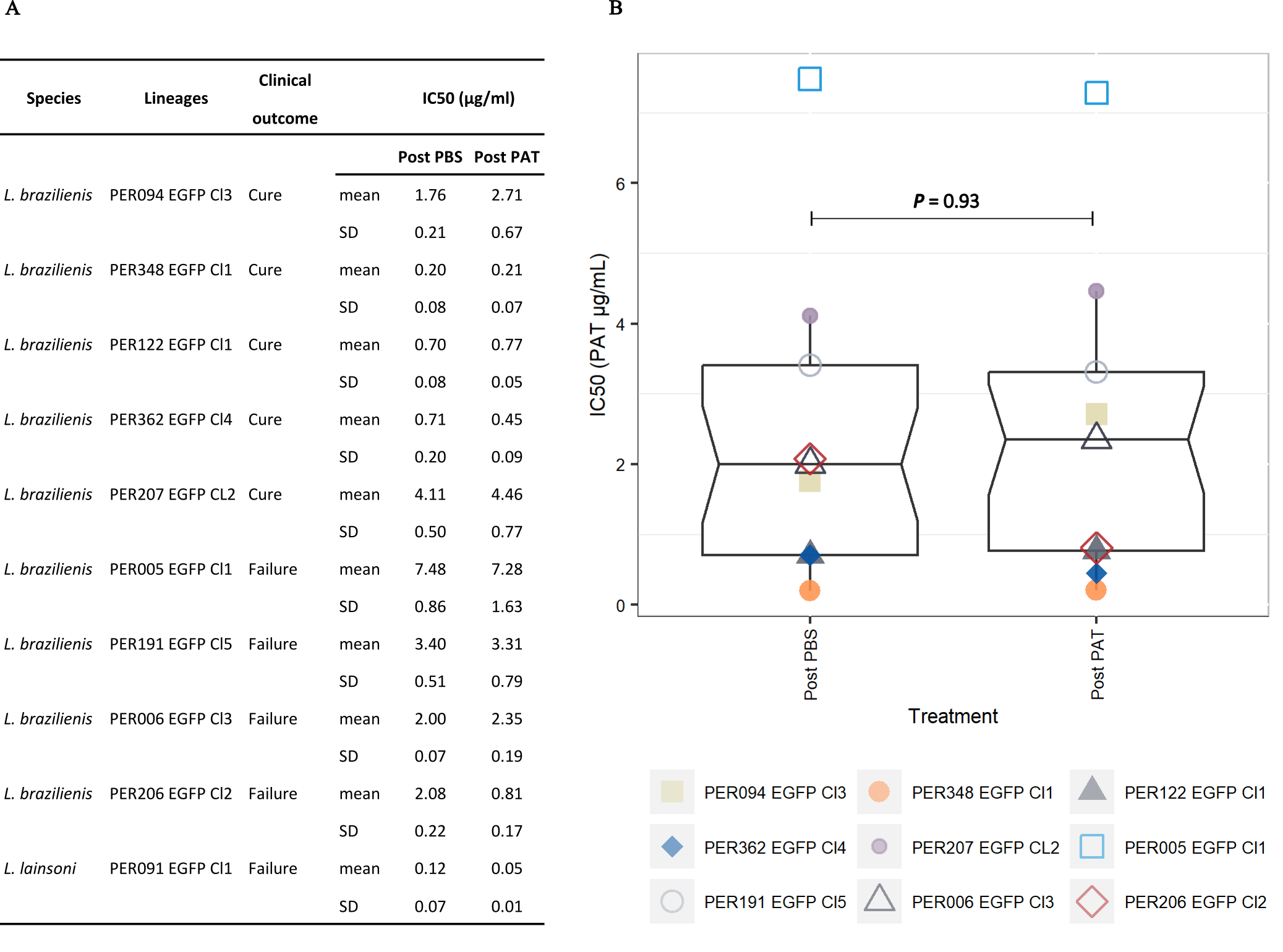

**Supplementary Figure 1** The overall susceptibility of *Leishmania* to PAT does not change in lineages after exposure to the drug compared to lineages without exposure. Promastigotes of each lineage were split into two groups one that was exposed to PAT (9 µg/mL) over 48 hrs and a control group treated with PBS. The survivors after PAT treatment were sub-cultured in fresh medium without drug pressure and kept in culture until population size allowed the resazurin test. The controls Post-PBS were also maintained in culture simultaneously until the evaluation of the IC50. **A.** The table shows the mean IC50 for three biological replicates as measured with the resazurin test after 24 hrs of exposure to PAT. The *L. lainsoni* strain PER091 EGFP Cl1 was used as a reference for high susceptibility to PAT. **B.** Box plot showing the IC50s of *L. braziliensis* lineages Post PBS and Post PAT. There was no significant difference between the median of both groups (Mann–Whitney U test, P= 0.93). IC, inhibitory concentration; PAT, Potassium antimonyl tartrate.

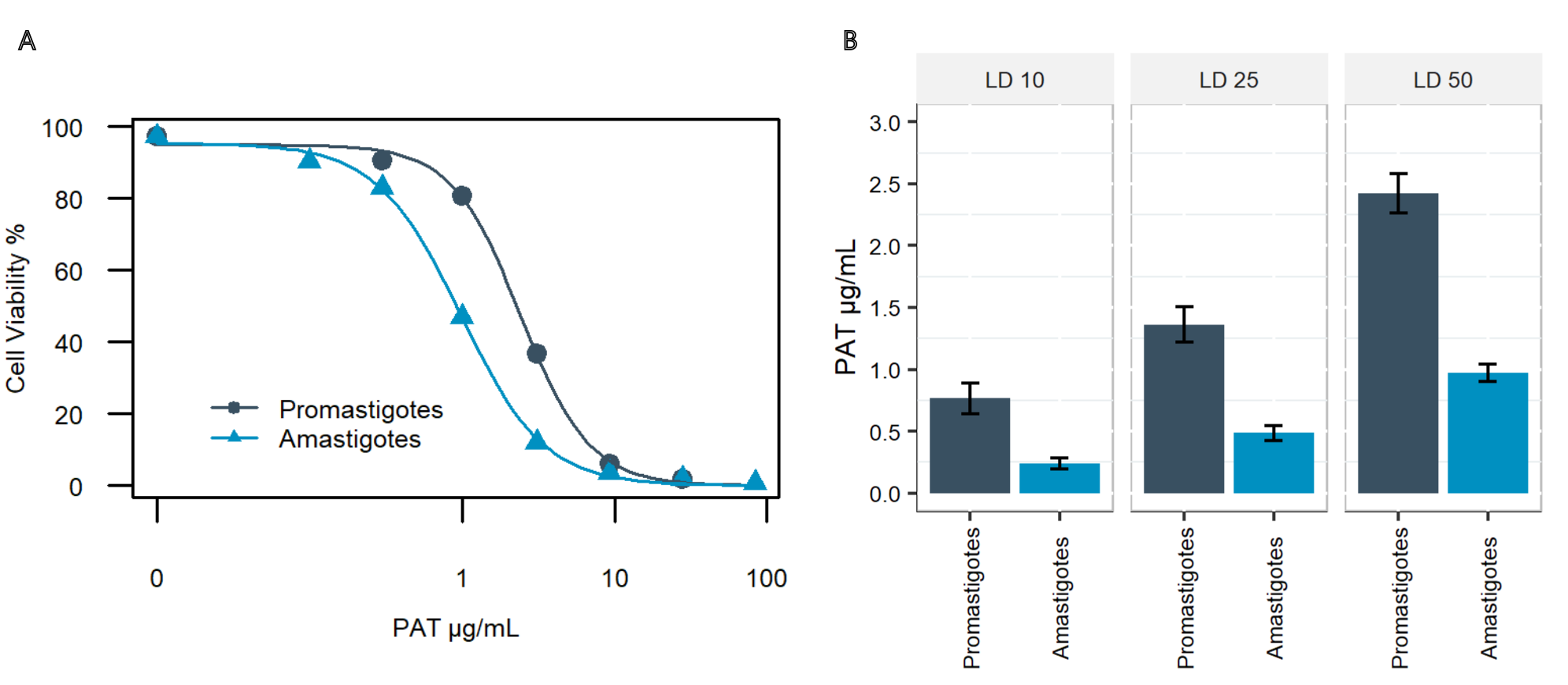

**Supplementary Figure 2.** Estimation of the LD10, LD25 and LD50 in response to PAT in an *L. lainsoni* strain after 48 hrs of exposure to PAT. We used the strain PER091 EGFP Cl1 which was previously characterized as highly susceptible to PAT in both promastigotes and amastigotes as a control for our experimental setup to calculate the LDs in amastigotes from *L. braziliensis*. The results indicate that PER091 EGFP Cl1 amastigotes are highly susceptible to PAT (LD_25_= 0.49 µg/mL) under the same experimental conditions compared to *L. braziliensis* (LD_25_ median =46.7 µg/mL). This indicates that the increased tolerance to PAT observed in amastigotes from *L. braziliensis* compared to their respective promastigotes is a biological trait and not a technical artefact. LD, Lethal dose. PAT; Potassium antimonyl tartrate.

**Supplementary data** (attached).

Genomic analyses. (i) Overall table summarizing for each strain the number and types of SNPs and indels observed before (Post-PBS) and after (Post-PAT) drug pressure; (ii) list of high impact indels; (iii) list of high impact SNPs.
